# A novel class of allosteric glucosylceramidase beta 1 correctors that reduce cellular stress and enhance lysosomal function

**DOI:** 10.1101/2024.11.15.623779

**Authors:** Ilaria Fregno, Natalia Pérez-Carmona, Mikhail Rudinskiy, Tatiana Soldà, Timothy J. Bergmann, Ana Ruano, Aida Delgado, Elena Cubero, Manolo Bellotto, Ana María García-Collazo, Maurizio Molinari

## Abstract

Mutations in glucosylceramidase beta 1 (GCase) disrupt the protein’s conformational maturation in the endoplasmic reticulum (ER) and hinder its transport to the lysosome. The intralysosomal accumulation of glucocerebrosides, which are substrates of the GCase enzyme, impairs lysosomal function and is linked to Gaucher disease (GD). GCase mutations also increase the risk of Parkinson’s disease (PD) and Dementia with Lewy Bodies. We used Site-directed Enzyme Enhancement Therapy (SEE-Tx®) technology to design two structurally targeted allosteric regulators (STARs) of GCase. Administration of GT-02287 and GT-02329 to cultured GD patient-derived primary human fibroblasts enhances folding and protects the two most common disease-causing GCase variants, GCase_Asn370Ser_ and GCase_Leu444Pro_, from proteasomal degradation. Mechanistically, these treatments facilitate the lysosomal delivery of enzymatically active forms of mutant GCase, leading to improved lysosomal function and reduced cellular stress in GD patient-derived fibroblasts. The findings suggest that the allosteric pharmacologic regulators GT-02287 and GT-02329 hold promise for further development as potential therapeutic agents for GCase-related disorders, including GD, PD and Dementia with Lewy Bodies.

## Introduction

Glucosylceramidase beta 1 (GCase) is a lysosome-resident protein that cleaves the beta-glucosidic linkage of glycosylceramides, generating glucose and free ceramides. GCase mutations may hamper folding of the polypeptide chain and result in unwanted clearance of the protein and/or reduced transport from the endoplasmic reticulum (ER) to the lysosomal compartment. Defective lysosomal delivery of functional GCase causes toxic intralysosomal accumulation of GCase substrates. Mutations in GCase are linked to Gaucher disease (GD), a lysosomal storage disorder (LSD), and are also a risk factor for Parkinson’s disease (PD) and Dementia with Lewy Bodies (Do, McKinney *et al*., 2019). The most common mutation in GCase is the substitution of the asparagine residue (N in the single letter code) at position 370—or position 409 if the 39 residues of the leader peptide are included in the amino acid numbering—with a serine residue (S). The GCase_Asn370Ser_ mutation is associated with type I GD, which typically presents a mild phenotype. Another common mutation is the replacement of the leucine residue (L) at position 444—or 483 if including the leader peptide—with a proline (P). This mutation often results in a more severe outcome, including the activation of unfolded protein responses (UPR) in both GD patients and a Drosophila model of the disease (Maor, Rencus-Lazar *et al*., 2013).

In this study, we used the HaloTag (HT) reporter approach developed in our lab (Rudinskiy, Bergmann *et al*., 2022, Rudinskiy and Molinari, 2022, Rudinskiy, Pons-Vizcarra *et al*., 2023), combined with an automated, deep learning method for segmenting and classifying fluorescence images, known as LysoQuant (Morone, Marazza *et al*., 2020). This approach captures cargo delivery within endolysosomes to monitor the impact of the disease-causing Asn370Ser and Leu444Pro mutations on the lysosomal delivery of GCase and the induction of cellular stresses. Moreover, we assessed the beneficial effects of structurally targeted allosteric regulators (STARs) of GCase, developed using the innovative drug discovery platform SEE-Tx^®^ (Barroso, Puchwein-Schwepcke *et al*., 2024, Cubero, Ruano *et al*., 2024, Rudinskiy, Pons-Vizcarra *et al*., 2023).

## Materials and methods

### Binding studies by surface plasmon resonance (SPR)

Human full-length wild-type GCase protein (Cerezyme, Genzyme, Naarden, NL) was immobilized on an SPR CM5 sensor (GE Healthcare, #29149603) using standard amino coupling with a high protein concentration of 100 µg/mL. A nine-point, 2-fold serial dilution, starting from 100 µmol/L GT-02287 and GT-02329 (prepared from a 10 mmol/L stock solution in DMSO), was measured by SPR at two pH conditions: pH 7.4 in 10 mmol/L HEPES, 5 mmol/L EDTA, 150 mmol/L NaCl and 0.01% Tween-20; and pH 5.0 in 20 mmol/L Na phosphate, 2.7 mmol/L KCl, 137 mmol/L NaCl, 5 mmol/L tartrate, 0.01% Tween-20. Reference channels on the SPR sensor included empty, activated, and deactivated parallel channels. Raw SPR signals from the active channel (with immobilized GCase protein) were double-referenced by subtracting signals from the reference channel (empty sensor surface) and the running buffer, with corrections for DMSO signal mismatch between the sample and the buffer. Binding affinity values were extracted by fitting the plotted SPR data with a four-parameter logarithmic dose-response equation using GraphPad Prism version 10.2.3. To assess the mode of action of the analyzed compounds, competition experiments were conducted with isofagomine (IFG), an inhibitory reference compound, present in the running buffer at saturating concentrations under both pH conditions (pH 7.4 and 5.0). These competition experiments were performed as single measurements and repeated twice.

### GCase biochemical and competition assay in wild-type lysates

Lysates were prepared from wild-type fibroblasts using a lysis buffer containing 0.9 % NaCl and 0.01 %Triton-X100. The protein concentration was determined using a BCA protein assay kit. To inhibit 50% of GCase activity, cell lysates were pre-incubated with 40 µM conduritol-β-epoxide (CBE) for 15 minutes. Subsequently, GT-02287 or GT-02329 was added for another 15 minutes. The assay was conducted in a reaction buffer containing 5 mM 4-methylumbelliferyl-beta-D-glucopyranoside in 0.1 M citrate-phosphate buffer at pH 5.6. Reactions were incubated at 37°C for one hour. To stop the reaction, 140 µL of 100 mM Glycine-NaOH buffer at pH 10.7 was added. The released 4-MU was measured using a Glomax® Discover microplate reader with excitation at 340 nm and emission at 460 nm.

### Cell culture, transient transfection, and use of compounds

Mouse embryonic fibroblasts (MEF) and Human Embryonic Kidney 293 (HEK293) cell lines were cultured at 37°C and 5% CO_2_ in Dulbecco’s Modified Eagle Medium (DMEM) with high glucose (GlutaMAX™, Gibco) supplemented with 10% Fetal bovine serum (FBS, Gibco). Fibroblasts from GD patients were obtained from the Coriell Institute for Medical Research and the Telethon Network of Genetic Biobanks. The specific genotypes used were as follows: p.N370S/ins (GD type I, Coriell GM00372), p.L444P/p.L444P (GD type II, Coriell GM08760), p.L444P/p.L444P (GD type III, Telethon 20526), p.L444P/p.L444P (GD type I, Coriell GM10915), p.L444P/p.F213I (GD type III, Telethon 21142), p.L444P/p.R496C (GD type III, Telethon 20624), and p.N188S/p.S107L (GD type III, Telethon 20843). Healthy fibroblasts (Coriell GM03377) were also included.

For enzyme enhancement and substrate depletion studies, fibroblasts were cultured in DMEM supplemented with 10% heat-inactivated FBS and 1% penicillin-streptomycin at 37°C under 5% CO_2_. Patient fibroblasts used in other studies were cultured in DMEM-GlutaMAX™ with 15% non-inactivated FBS under similar conditions.

Transient transfections were conducted using JetPrime (Polypus) in DMEM 10% FBS supplemented with non-essential amino acids (NEAA, Gibco) following the manufacturer’s protocol. Pharmacological chaperones from Gain Therapeutics (GT), GT-02287 and GT-02329, were dissolved in dimethylsulfoxide (DMSO, Sigma) and used at concentrations specified in the figure legends. The GCase inhibitor, conduritol-β-epoxide (CBE), was acquired from Merck Millipore (Catalogue No: 234599). The proteasome inhibitor PS341 (Bortezomib, LubioScience) was used at a final concentration of 10 µM.

### Plasmids and cloning

Plasmids encoding for GCase-HA (human influenza hemagglutinin), GCase-Halo Asn370Ser and Leu444Pro mutant polypeptides within the pcDNA3.1(+) backbone, flanked by HindIII and XhoI restriction sites were synthesized by GenScript. The transfer of HA and HaloTag tags between various plasmids was achieved through restriction digestion of NotI and XhoI sites flanking the tags. Ligation was conducted using T4 DNA ligase (NEB) at a 3:1 insert to vector ratio, following the manufacturer’s protocol. Plasmids were subsequently amplified and isolated from JM109 bacteria (Promega) using the GenElute™ HP plasmid MidiPrep kit (Sigma).

### Confocal laser scanning microscopy (CLSM)

MEFs were seeded on alcian blue-treated (Sigma) glass coverslips (VWR) and transiently transfected using JetPrime (Polypus) in DMEM 10% FBS supplemented with non-essential amino acids (NEAA; Gibco) according to the manufacturer’s protocol. Thirty-two hours after transfection, bafilomycin A1 (BafA1; Calbiochem) was added to the cell medium at the final concentration of 50 nM for 17 h. Cells expressing HaloTag-fusion proteins were supplemented with 100 nM tetramethylrhodamine (TMR) HaloTag ligand (Promega). After treatment, MEFs were fixed at room temperature (RT) for 20 min in 3.7% formaldehyde (FA, v/v) in phosphate buffered saline (PBS). Coverslips were incubated for 20 min in permeabilization solution (PS, 10 mM HEPES, 15 mM glycine, 10% goat serum (v/v), 0.05% saponin (w/v)). Following permeabilization, primary antibodies diluted 1:100 in PS (unless otherwise specified in Table **S1**) were applied for 120 min, washed three times in PS, and then Alexa Fluor-conjugated secondary antibodies diluted 1:300 in PS were applied for 45 min. Cells were washed three times with PS and once with deionized water and mounted on a drop of Vectashield (Vector Laboratories) supplemented with 40,6-diamidino-2-phenylindole (DAPI). Coverslips were imaged using a Leica TCS SP5, microscope equipped with Leica HCX PL APO lambda blue 63.0 × 1.40 oil objective. Leica LAS X software was used for image acquisition, with excitation provided by 488-, 561-, and 633 nm lasers. Fluorescence light was collected within the ranges of 504-587 nm (AlexaFluor488), 557-663 nm (TMR) and 658-750 nm (AlexaFluor646), respectively, with a pinhole setting of 1 AU. The accumulation of GCase variants within LAMP1-positive endolysosomes was quantified with LysoQuant plugin within the FIJI/ImageJ software (Morone, Marazza *et al*., 2020, Schindelin, Arganda-Carreras *et al*., 2012). Image post-processing was performed with Adobe Photoshop.

### Sodium dodecyl sulfate polyacrylamide gel electrophoresis (SDS-PAGE), HaloTag cleavage assay, co-Immunoprecipitation and western blot (WB)

HEK293 cells or patient fibroblasts were washed with ice-cold 1xPBS containing 20 mM N-ethylmaleimide (NEM), then lysed with RIPA (1% Triton X-100 (v/v), 0.1% SDS (w/v), 0.5% sodium deoxycholate in HBS, pH 7.4, (w/v)) or 2% CHAPS (in HEPES-buffered saline [HBS], pH 7.4, (w/v)) buffer containing 20 mM NEM and protease inhibitors (1 mM phenylmethylsulfonyl fluoride (PMSF), 16.5 mM Chymostatin, 23.4 mM Leupeptin, 16.6 mM Antipain, 14.6 mM Pepstatin) for 20 min on ice. The lysate was centrifuged to extract the post-nuclear supernatant (PNS) for 10 min at 4°C, 10,600 g. The PNS was denatured and reduced by adding 100 mM dithiothreitol (DTT, Roche) and heating the sample for 5 min at 95°C. Boiled samples were subjected to 12% Acrylamide SDS-PAGE. Lysates of cells expressing GCase-HT variants labeled with fluorescent TMR Halo ligand (Promega) were imaged *in-gel* with Typhoon™ FLA 9500 fluorescent scanner (GE Healthcare) with a 532 nm laser.

The intensity of TMR bands was quantified using the ImageQuantTL software (Molecular Dynamics, GE Healthcare). Total protein content was quantified by staining the polyacrylamide gel with 0.25% Brilliant blue R 250 (Sigma) solution (50% MeOH (v/v), 10% acetic acid (v/v)) for 20 min at RT, and destaining in 20% MeOH (v/v), 7.5% acetic acid (v/v) solution. The protein bands were acquired on a Fusion FX7 system (Vilber) with transilluminator and quantified with FIJI/ImageJ software. For immunoprecipitation, the lysates of cells treated with DMSO or GT compounds were diluted with lysis buffer and incubated with the select bait antibodies and protein A-conjugated beads (1:10 w/v, swollen in PBS) at 4°C for 4 hours. The beads were centrifuged down and washed three times with 0.5% Triton X-100 (v/v) in HBS pH 7.4. After that, beads were denatured for 5 min at 95°C, and the liquid fraction was subjected to SDS-PAGE.

For detection of protein band by WB protein bands were transferred from polyacrylamide gel to polyvinylidene fluoride (PVDF) membrane using a TransBlot Turbo device (BioRad). PVDF membrane was blocked for 10 min with 10% blocking milk (BioRad, w/v) in Tris-buffered saline, 0.1% Tween 20 (v/v) (TBS-T), rinsed quickly in TBS-T and incubated with primary antibodies (**Table S1**) overnight at 4°C with shaking. After primary antibody washout with TBS-T, membranes were incubated with HRP-conjugated secondary antibody or protein A (**Table S1**) for 45 min at RT with shaking. Protein bands were detected on a Fusion FX7 chemiluminescence detection system (Vilber) using the WesternBright™ Quantum (Advansta) system following the manufacturer’s protocol. Protein bands were quantified with FIJI/ImageJ software.

### Radioactive metabolic labelling

Four days after the start of treatment with DMSO or GT compounds, patient fibroblasts were pulsed 4 h with 0.2 mCi [35S] methionine/cysteine mix and chased for 10 min with DMEM supplemented with 5 mM cold methionine and cysteine. Cells were solubilized in RIPA buffer, and radiolabeled proteins were revealed with Typhoon™ FLA 9500 fluorescent scanner (GE Healthcare). The intensity of radioactive bands was quantified using the ImageQuantTL software (Molecular Dynamics, GE Healthcare).

### RNA extraction and reverse transcription-quantitative polymerase chain reaction

Patient fibroblasts treated with DMSO or GT compounds were detached with trypsin and then centrifuged for 5 min at 450*g*. Total mRNA was extracted using the GenElute^TM^ mammalian total RNA Miniprep kit (Sigma) according to manufacturer’s protocol. The isolated mRNA was reverse transcribed into cDNA for quantitative polymerase chain reactions (qPCRs) using the BioTool™ 2x SYBR Green qPCR Master Mix (BioTool) as per the manufacturer’s instructions. The master mix was combined with qPCR primers, which were obtained from Microsynth AG and used at a final concentration of 10 µM, on the reading plate. qPCRs were conducted using the QuantStudio™ 3 Real-Time PCR System (Applied Biosystems). Data analysis was performed using QuantStudio™ Design & Analysis Software v1.5.5 (Applied Biosystems).

### GCase enzyme enhancement assay

Patient-derived fibroblasts were seeded at 5,000 cells per well in 96-well cell culture plates in DMEM supplemented with 10% FBS, 1% P/S (Thermo Fisher Scientific, Waltham, MA, USA) and incubated at 37°C, 5% CO2 overnight for cell attachment. Subsequently, cells were incubated with or without the specified compound at the indicated concentrations for 4 days. After incubation, cells were washed with PBS and incubated with 5 mM of 4-methylumbelliferyl-beta-D-glucopyranoside (Apollo Scientific) in 0.1 M acetate buffer pH 4 for 1 h at 37°C. The reaction was stopped by adding 200 µL of 100 mM Glycine-NaOH buffer, pH 10.7. The liberated, 4-methylumbelliferyl (4-MU) was measured on a Glomax® Discover microplate reader with excitation at 340 nm and emission at 460 nm.

### GlcCer quantification in patient-derived fibroblasts

Patient-derived fibroblasts were seeded at 750,000 cells in 75 cm^2^ flasks in DMEM supplemented with 10% FBS, 1% P/S. The following day, fibroblasts were treated with compounds at the indicated concentrations. On day 3 and day 7 of the treatment, supernatant was discarded and new media with compound was added. On day 10, supernatant from the cell culture was discarded and fibroblasts were trypsinized. Growth cell media was then added to the detached cells and collected in tubes. Next, the pellets were centrifuged in cool PBS and after discarding the supernatants, and the pellets were stored at -80°C.

GlcCer quantification was performed as follows: The pellets from the cell culture samples were transferred into Eppendorf tubes and rapidly weighed. The tubes were stored in an ice-water bath, and to each tube deionized water was pipetted in a 1:10 (weight/volume) ratio. Deuterium-labelled GlcCer-D5 was used as internal standard. The pellets were resuspended in water by use of an ultrasonic homogenizer VCX130 (Sonics & Materials, CT, USA) and 10 μl from each vial was pipetted for quantification of protein content. From the remaining volumes, 70 μl was pipetted into the separate vials and GlcCer was extracted from the resuspended cell debris using liquid-liquid extraction. The ultra-high performance liquid chromatography coupled with tandem mass spectrometry (UHPLC-MS/MS) system for quantification of glycosphingolipids included a Xevo TQ-S micro triple quadrupole mass spectrometer, an ACQUITY UPLC system consisting of a binary solvent manager, a sample manager operating at +8°C and equipped with a 10 μl loop, a high temperature column heater, and a solvent tray module (all from purchased from Waters Micromass, Milford, MA, USA). The mass spectrometer was supplied by nitrogen gas, generated by a Genius 3045 nitrogen/air generator (Peak Scientific Instruments, Inchinnan, Scotland, UK). The electrospray ionization (ESI) source operated in a positive mode with the following parameters: Capillary voltage 4500 V, sample cone voltage 30 V, source temperature 150°C, desolvation temperature 500°C, cone gas flow 0 L/hr, nebulizer gas flow 1000 L/hr. Acquisition parameters were: Centroid MRM scans of parent and daughter ions, 0.1 second dwell time. The divert valve was programmed as follows: 0.00 min waste, 1.30 min run, 3.2 min waste. The chiral separation of GlcCer from GalCer was achieved on a HALO HILIC column (2.1 x 150 mm, 2 μm particle size, purchased from Advanced Materials Technology, Wilmington, DE, USA) and maintained at +35°C. The mobile phase consisted of 95% ACN, 2.5% MeOH, and 2.5% water containing 5 mM ammonium formate and 0.5% formic acid; the flow rate was 0.240 ml/min. The protein content in 10 μl volumes of the resuspended samples was measured by Pierce Rapid Gold BCA protein assay (Thermo Fisher Scientific, USA) and an UV/VIS plate reader (Labrox, Finland). The results for GlcCer content in the pellets were presented as ng/mg protein. Data analysis was performed using GraphPad Prism 5.0 Software. All values show the mean +/- SD (standard deviation) from 3 wells per condition. Statistical differences *vs.* untreated GCase_Leu444Pro_ were determined by one-way analysis of variance (ANOVA) followed by a Dunnett’s Multiple Comparison Test. Asterisks indicate statistically significant values from untreated GCase_Leu444Pro_ **** p≤0.0001 and *** p≤0.001.

### Statistical analyses

For figures **1**‒**4**: Data analysis was performed using GraphPad Prism 5.0 Software. For figures **5****‒8**: Statistical comparisons and graphical plots were performed in GraphPad Prism 10 (GraphPad Software Inc.). An ordinary one-way ANOVA Dunnett’s multiple comparisons test was used to assess statistical significance (for **Figs. 4A** and **4B**). An adjusted p-value < 0.05 (for one-way ANOVA with Dunnett’s multiple comparison test) was considered statistically significant: *** p<0.001 and **** p<0.0001. For figures **5C**, **5J, 6B, 7D** and **7G**, an unpaired t-test was used to assess statistical significance. An exact p-value <0.05 was considered significant; ns is not significant, *p<0.05, **p<0.01, ***p<0.001 and ****p<0.0001. All replicates represent biological replicates, and for all statistical comparisons the number of repetitions and the number of cells analyzed (n, for LysoQuant analyses) are indicated in the figure legends.

**Figure 1.**
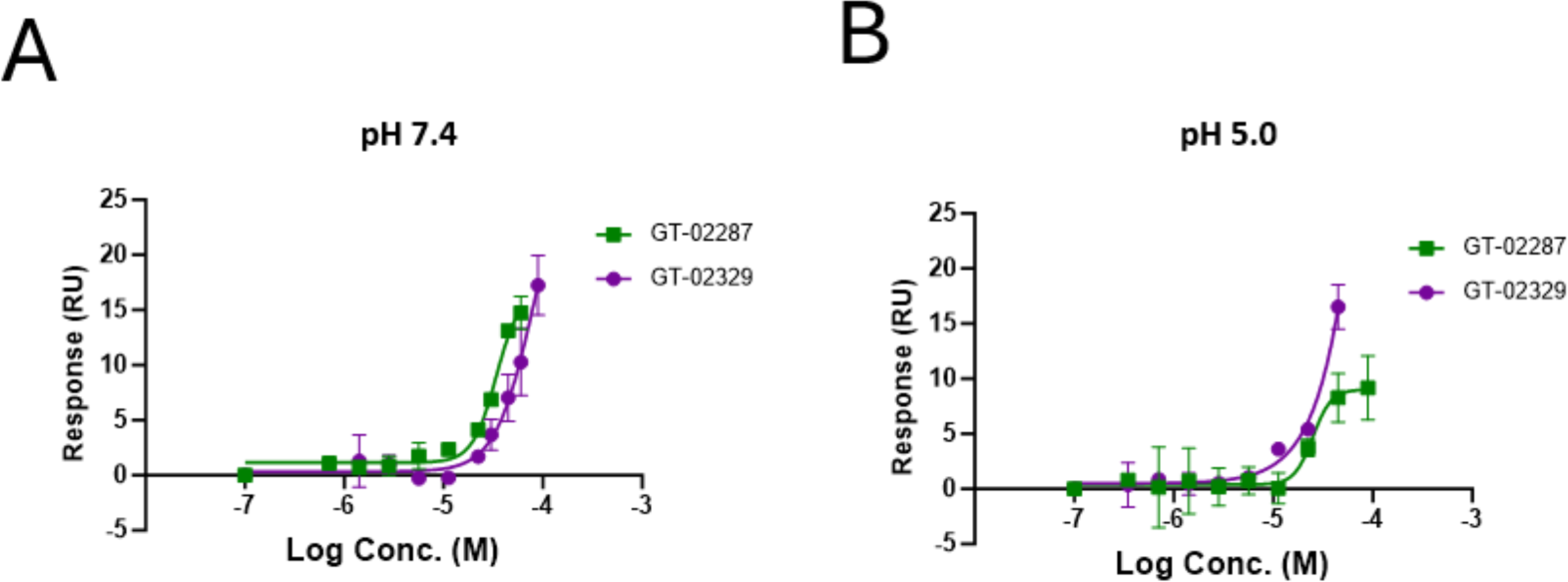
GCase binding confirmation. Surface plasmon resonance direct binding measured for compounds GT-02287 and GT-02329 at (A) pH 7.4 and (B) pH 5.0. All values show the mean +/- SD (standard deviation) in duplicates.

## Results

### GT-02287 and GT-02329 bind to the GCase protein in an allosteric pocket

SEE-Tx^®^ technology (Alonso, Garcia *et al*., Patent 2012, Alvarez-Garcia and Barril, 2014, Alvarez-Garcia, Schmidtke *et al*., 2022) utilized the high-resolution structure of native human GCase (PDB: 2F3V) (Shaaltiel, Bartfeld *et al*., 2007) to identify an allosteric binding site and predict its druggability. This process guided subsequent steps: key binding hotspots informed a high-throughput virtual screening assay (Ruiz-Carmona, Alvarez-Garcia *et al*., 2014), leading to the identification of novel virtual hits that were purchased and tested for interaction with the GCase protein. Experimentally validated hits served as starting points for a medicinal chemistry program, resulting in the selection of GT-02287 and GT-02329. Notably, chemical structures remain undisclosed due to the sensitive nature of proprietary research in the drug development phase.

The direct binding of GT-02287 and GT-02329 to the GCase target protein was confirmed by SPR. SPR provides real-time measurements of molecular interaction kinetics and affinity, making it an optimal technique for this investigation (Schuck, 1997). These experiments were conducted at pH 7.4 and 5.0, mimicking the ER and the luminal environment of acidic organelles, respectively (Patching, 2014). GT-02287 and GT-02329 showed a dose-dependent binding to GCase protein at neutral and acidic conditions, with comparable binding affinities at pH 7.4 and 5.0 (**Figs. 1A, 1B**).

SPR competition experiments with IFG (Park, Chen *et al*., Shire Human Genetic Therapies, Inc. Patent 2019), a competitive inhibitor of GCase, confirmed that GT-02287 and GT-02329 bind to an allosteric site and do not block the active site on GCase. IFG binds to the active site of the GCase protein and acts as a pharmacological chaperone, stabilizing mutant enzymes to improve their function. Dissociation constant (K_D_) values obtained with and without IFG were comparable at pH 7.4 and 5.0, indicating that these compounds bind to a pocket distinct from the active site occupied by IFG (**Table 1**; see also supporting information).

**Table 1.** SPR direct binding affinities measured at pH 7.4 and 5.0, and competition data with isofagomine (IFG). K_D_, dissociation constant.

| Compound | pH 7.4 |  | pH 5.0 |  | Competition with IFG |
| --- | --- | --- | --- | --- | --- |
|  | K<sub>D</sub> (μM) | K<sub>D</sub> (μM) + IFG | K<sub>D</sub> (μM) | K<sub>D</sub> (μM) + IFG |  |
| **IFG (control)** | 0,03 – 0,06 | - | 0,10 – 0,11 | - | - |
| **GT-02287** | 17 - 24 | 27 - 30 | 50 - 150 | 55 - 64 | No |
| **GT-02329** | > 90 | > 90 | 39 - 40 | 30 - 31 | No |

### GT-02287 and GT-02329 enhance GCase activity in fibroblast lysates by binding to a site on the enzyme different from the active site

The effect of GT-02287 and GT-02329 on the enzymatic activity of GCase was assessed in wild-type fibroblast extracts at pH 5.6. Both compounds enhance GCase activity at concentrations above 10 µM, as evidenced using the substrate 4-methylumbelliferyl-β-D-glucopyranoside (**Figs. 2B and 2C**). These results support the notion that the compounds bind to a site that is different from the catalytic site. Otherwise, they would be expected to exhibit inhibitory behavior, as observed with known inhibitors such as IFG.

**Figure 2.**
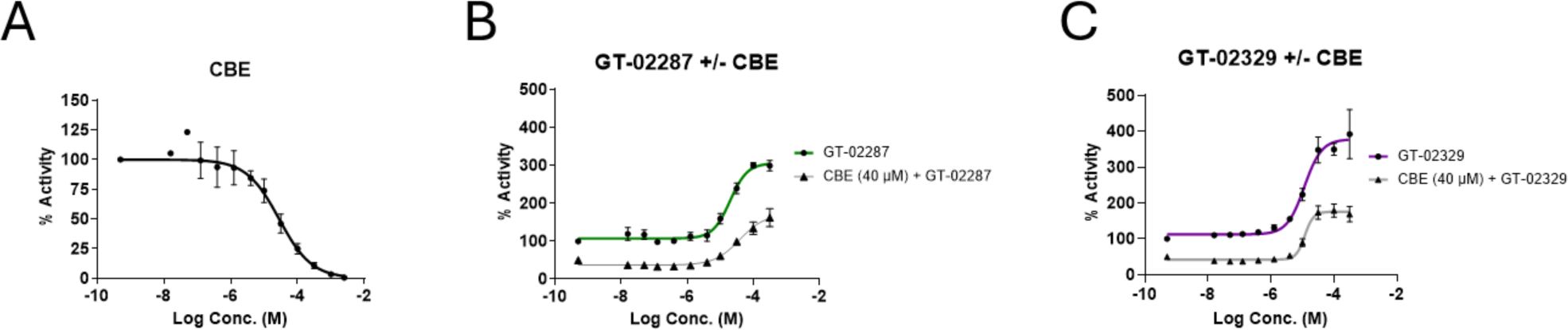
Competitive inhibition assay with CBE in lysates. Wild-type fibroblast lysates were incubated with (A) CBE black), (B) GT-02287 (green) or (C) GT-02329 (purple) or for 15 minutes at the indicated doses. GCase activity was then measured using the 4-methylumbelliferyl β-D-glucopyranoside substrate. Where indicated, an additional preincubation with CBE at 40 µM (**C**, **D** in grey) was performed to inhibit 50% of GCase activity. All values show the mean +/- SD (standard deviation) from 3 wells per condition. CBE, conduritol-β-epoxide.

CBE, a potent, selective irreversible competitive inhibitor, covalently binds GCase in the active site with a IC_50_ of 28.19 µM, as shown in **Fig. 2A** (Premkumar, Sawkar *et al*., 2005). When the activity of GT-02287 and GT-02329 was assessed in the presence of CBE at a concentration that irreversibly inhibits 50% of GCase activity, the effect of the GT compounds on GCase enzymatic activity was reduced by 50% (**Figs. 2B and 2C**, respectively). This result demonstrates that the enhancement of GCase activity is mediated by binding to GCase, showing a clear relationship between GT-02287 and GT-02329’s activity enhancement and the availability of functional GCase in patient fibroblast extracts.

### GT-02287 and GT-02329 enhance GCase enzyme activity in primary human fibroblasts and from GD patients

The ability of GT-02287 and GT-02329 to enhance the activity of GCase proteins with disease-causing mutations was assessed in primary human fibroblasts from a healthy donor (**Figs. 3A and 3A’**) and from six patients affected by GD type I, GD type II and GD type III (**Figs. 3B‒3G and 3B’‒3G’**), resulting from various mutations in the GBA1 gene. Briefly, fibroblasts were seeded, and the following day, the compounds were added. After a 4-day incubation, GCase activity was measured using the fluorogenic substrate 4-methylumbelliferyl-β-D-glucopyranoside. In all cases, incubation of the primary human fibroblasts with GT-02287 and GT-02329 resulted in a dose-dependent increase in GCase activity.

**Figure 3.**
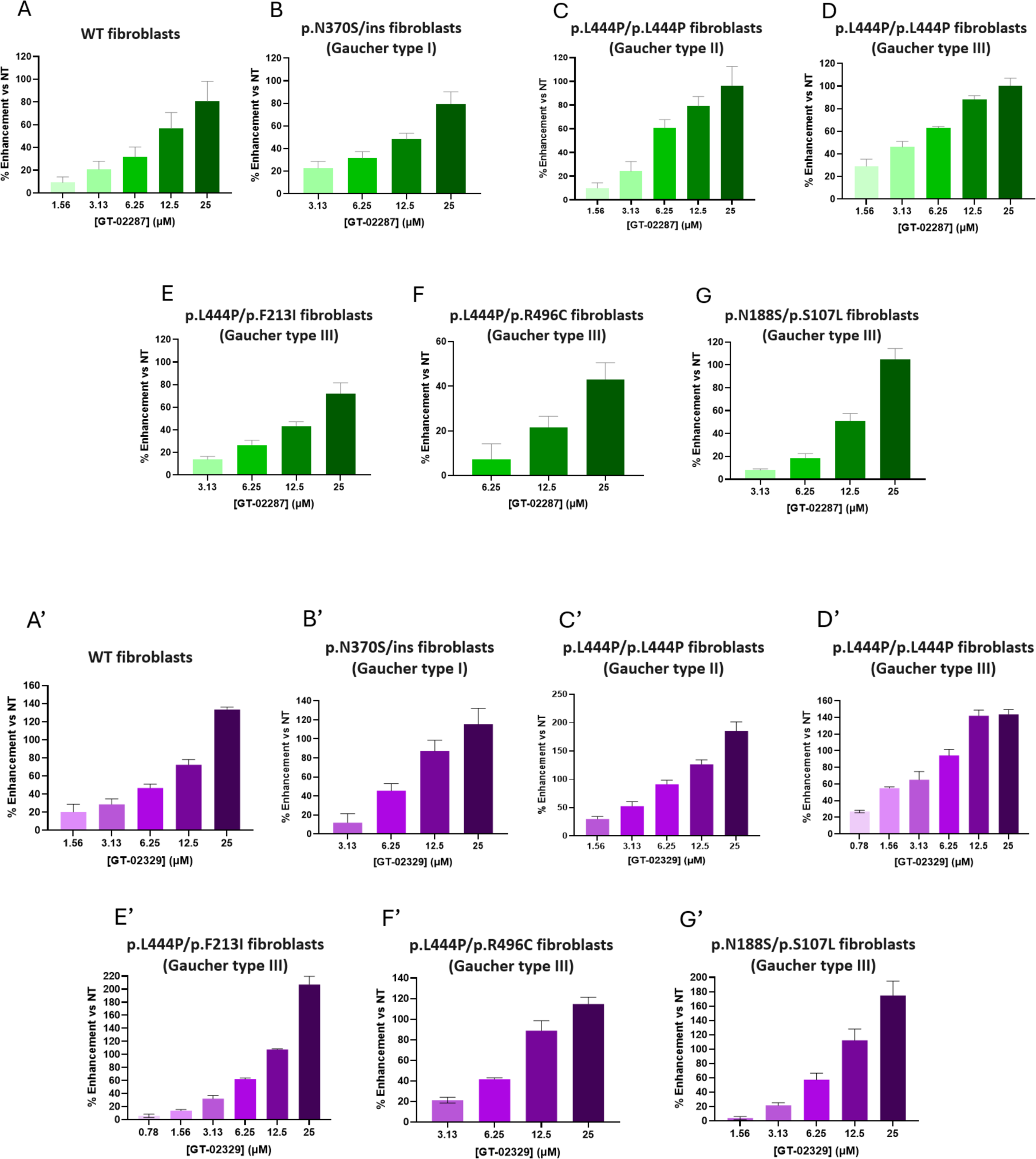
Cellular GCase enhancing activity of GT-02287 and GT-02329. Patient-derived fibroblasts were treated with GT-02287 (**A‒G**, green) or GT-02329 (**A’**‒**G’**, purple) at the specified doses for 4 days. GCase activity was measured using the 4-methylumbelliferyl β-D-glucopyranoside substrate. The dose-response effect is shown as the percentage increase in activity relative to untreated cells. Mean +/- standard deviation (SD) from at least three wells per condition are shown.

### GT-02287 and GT-02329 treatment reduces endogenous substrate accumulation in primary human fibroblasts

Reduced GCase activity in patient-derived p.L444P/p.L444P primary fibroblasts leads to an accumulation of glucosylceramide (GlcCer), a primary physiological substrate of GCase (**Fig. 4**). To determine whether enhancing GCase activity in patient cells by treating with GT-02287 or GT-02329 reduces the intracellular accumulation of GlcCer, fibroblasts were treated for 10-days, starting one day after seeding, after which, cells were harvested and processed as described in the methods section. Quantification by UHPLC-MS/MS revealed that treatment of patient fibroblasts with increasing concentrations of GT-02287 (**Fig. 4A**) or GT-02329 (**Fig. 4B**) effectively mitigates GlcCer accumulation.

**Figure 4.**
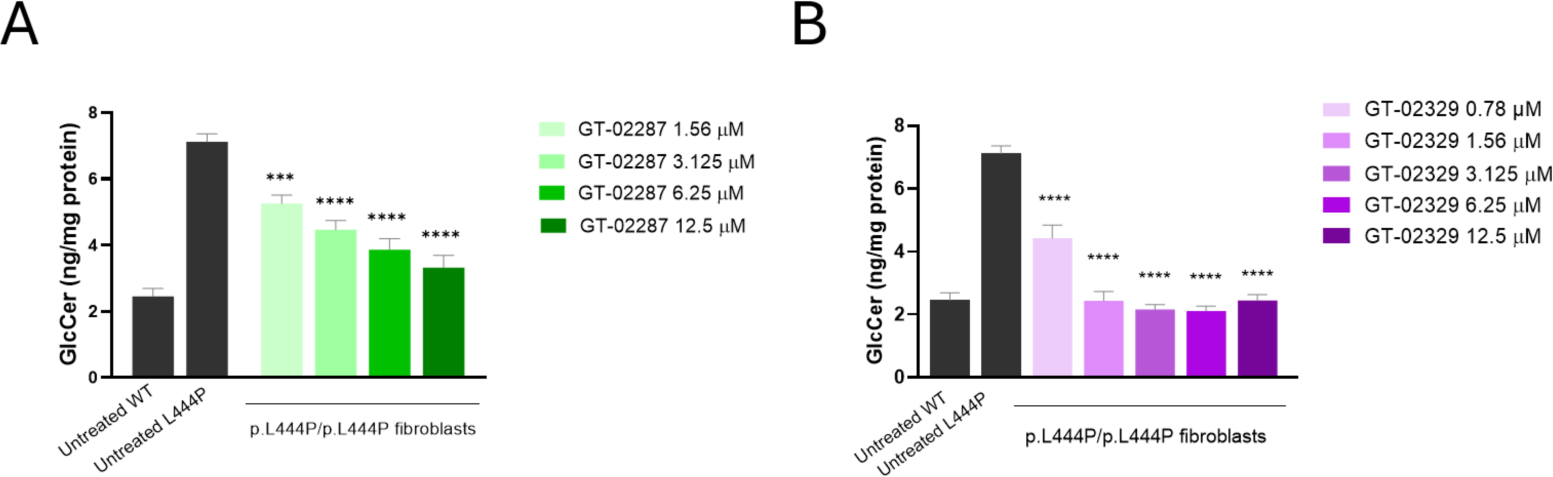
GlcCer substrate depletion following treatment with GT-02287 and GT-02329. p.L444P/p.L444P fibroblasts (Coriell GM08760) were treated with GT-02287 (**A**) or GT-02329 (**B**) at the specified doses for 10 days. GlcCer levels were quantified by ultra-high performance liquid chromatography coupled with tandem mass spectrometry (UHPLC-MS/MS). The mean +/- standard deviation (SD) is shown from 3 wells per condition. A one-way analysis of variance (ANOVA) followed by Dunnett’s Multiple Comparison Test was performed. Asterisks indicate statistically significant values from untreated GCase_Leu444Pro_ **** p≤0.0001, *** p≤0.001.

### HaloTagged chimeras to monitor defective lysosomal transport of disease-causing GCase mutations by CLSM

To quantitatively assess the performance and the mode-of-action of GT-2287 and GT-2329, we developed a biochemical protocol. This protocol has been instrumental in evaluating STARS that enhance the activity of the galactosidase beta 1 (GLB1) in GLB1-related LSDs, including GM1 gangliosidosis (Rudinskiy, Bergmann *et al*., 2022, Rudinskiy and Molinari, 2022, Rudinskiy, Pons-Vizcarra *et al*., 2023).

Briefly, GCase and two disease-causing variants (GCase_Asn370Ser_ and GCase_Leu444Pro_) were tagged with a HaloTag polypeptide to create GCase-HaloTag (GCase-HT) chimeras. HaloTag is an engineered bacterial dehalogenase, whose active site covalently binds cell-permeable haloalkanes modified with fluorescent ligands (England, Luo *et al*., 2015, Los, Encell *et al*., 2008). Chimeric polypeptides modified with HaloTag can be fluorescently labeled by adding cell-permeable fluorescent Halo ligands, such as tetramethylrhodamine (TMR), to the cell culture media. The intracellular distribution of HaloTagged proteins can be assessed by CLSM.

Given that HaloTag (297 residues) is much larger than conventional tags such as the HA-tag (9 residues) (Kimple, Brill *et al*., 2013), we first checked whether HaloTag perturbs the lysosomal delivery of GCase, the mature form of which is composed of 497 residues (**Fig. 5A**). GCase, GCase_Asn370Ser_ and GCase_Leu444Pro_ were tagged at their C-terminus with HA or HaloTag epitopes and transiently expressed in MEFs. The delivery of the GCase variants, immunostained with anti-HA antibody (red) in lysosomes marked with an anti-LAMP1 antibody (green), was assessed by CLSM (**Fig. 5B**) and quantified by Lysoquant to monitor the accumulation of the HA-tagged GCase in lysosomes inactivated by 50 nM BafA1 (Morone, Marazza *et al*., 2020) (**Fig. 5C**). Micrographs show that GCase-HA (**Fig. 5B**, upper panels) and GCase_Asn370Ser_-HA (**Fig. 5B**, middle panels) are efficiently transported to LAMP1-positive lysosomes (LysoQuant quantifications, **Fig. 5C**). In contrast, transport of the GCase_Leu444Pro_-HA mutant is highly defective (**Figs. 5B** lower panels and **5C**). This correlates with the GD phenotype, which is mild for GCase_Asn370Ser_ and severe for GCase_Leu444Pro_ patients (Do, McKinney *et al*., 2019).

**Figure 5.**
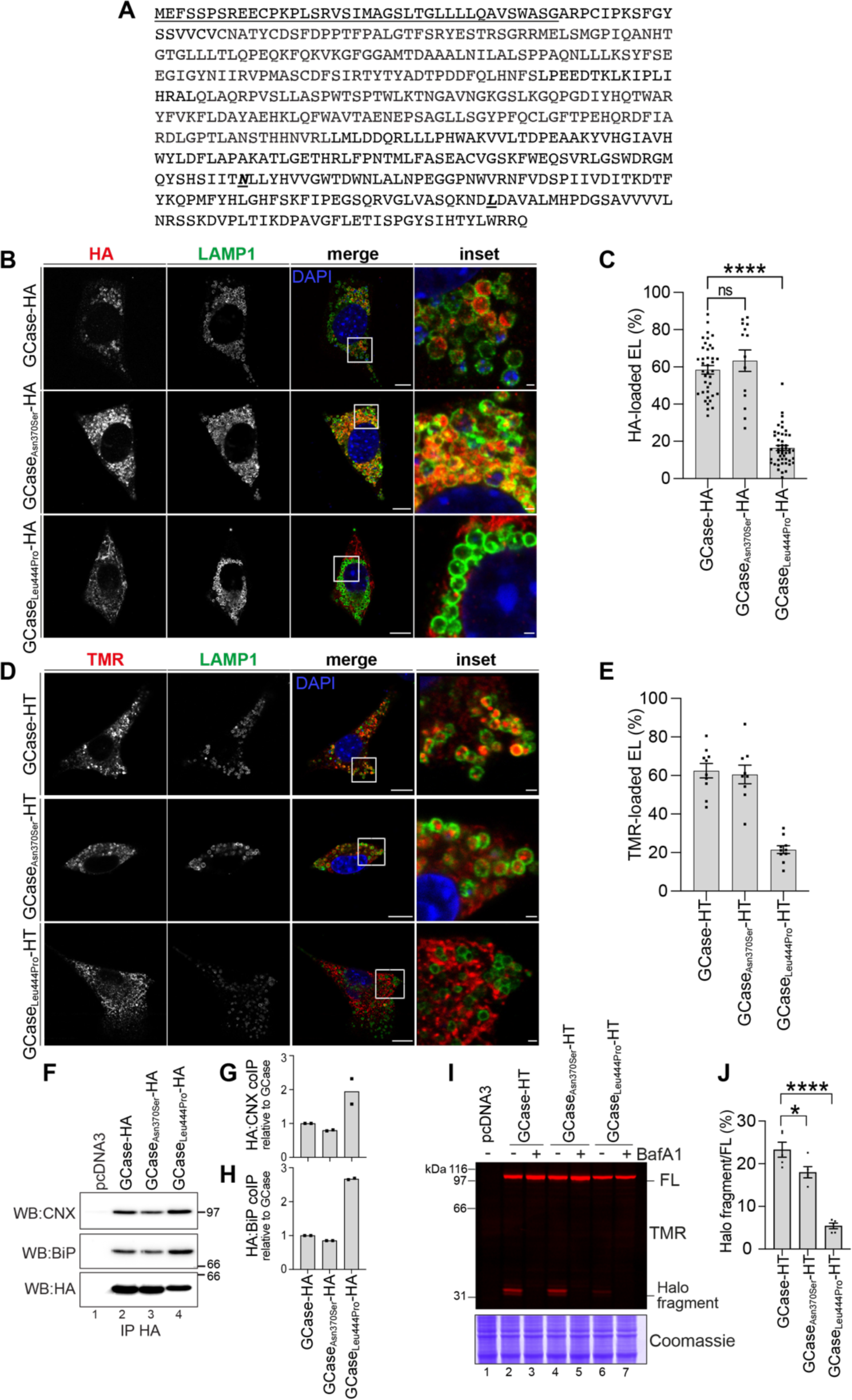
Lysosomal delivery of wild type and disease-linked GCase variants. **(A)** Sequence of GCase. The signal sequence is underlined. The sites of Asn_370_Ser and Leu_444_Pro mutations in the disease-causing gene products are shown in bold italics. (**B)** Lysosomal delivery of HA-tagged GCase variants. Representative CLSM images of MEF cells expressing GCase variants for 48 h and exposed for 17h to 50 nM bafilomycin A1 (BafA1). Immunostaining with anti-HA (red) anti-LAMP1 antibody (endolysosomes, green). Nuclei (blue) are stained with DAPI. Scale bars 10 µm (merge) and 1 µm (inset). (**C)** Quantification of (**B)** as a percentage of LAMP1-positive lysosomes containing GCase variants. Images analyzed with LysoQuant (Morone, Marazza *et al*., 2020). Mean -/+ SEM is shown, n=3 independent experiments. Statistical analysis: One-way ANOVA followed by Dunnett’s multiple comparison test, non-significant (ns) p>0.05, ****p<0.0001. (**D)** Same as (**B)** for Halotagged (HT) GCase variants. MEF cells expressing GCase variants for 48 h were exposed 17h to 50 nM BafA1 and 100 nM TMR. Staining for GCase variants with TMR (red) and lysosomes with anti-LAMP1 antibody (green). Scale bars 10 µm (merge) and 1 µm (inset). (**E)** Quantification of (**D)** as a percentage of LAMP1-positive lysosomes containing the TMR signal within their lumen. Images analyzed with LysoQuant (Morone, Marazza *et al*., 2020). Mean -/+ SEM is shown, n=2 independent experiments. (**F)** Co-immunoprecipitation of GCase-HA variants (WB: HA, lower panel and CNX, upper panel) and BiP (middle panel) in HEK293 cells. (**G)** Quantification of CNX association with the three GCase variants. (**H)** Same as **(G)** for BiP association. Mean is shown, n=2 independent experiments. (**I)** Fluorescent gel showing the TMR signal of GCase-HT variants expressed for 4 days in HEK293 cells in the presence or absence of 50 nM BafA1 for the last 17h. NT, non-transfected; FL, full-length GCase-HT. (**J)** Quantification of (**I)**. Mean -/+ SEM is shown, n=5 independent experiments. Statistical analysis: One-way ANOVA followed by Dunnett’s multiple comparison test, *p<0.05, ****p<0.0001. GCase glucocerebrosidase; HA hemagglutinin tag; CLSM, confocal laser scanning microscopy; MEF, mouse embryonic fibroblasts; BafA1, bafilomycin A1; LAMP1, lysosomal-associated membrane protein 1; DAPI, 4’,6-diamidino-2-phenylindole; EL, endolysosomes; SEM, standard error of the mean; HT, HaloTag; TMR, tetramethylrhodamine; CNX, calnexin; BiP, binding immunoglobulin protein; HEK293, human embryonic kidney 293 cells; WB, western blot; NT, non-transfected; FL, full length.

To assess intracellular trafficking of the same GCase variants when tagged with HaloTag, MEF cells were transiently transfected with the expression plasmids and the media supplemented with 100 nM TMR to label GCase-HaloTag variants. Their delivery to the LAMP1-positive compartment was assessed with CLSM (**Fig. 5D**) and quantified with LysoQuant (**Fig. 5E**). Results mirror those with HA-tagged polypeptides, i.e., GCase-HT and GCase_Asn370Ser_-HT are efficiently transported to LAMP1-positive lysosomes while GCase_Leu444Pro_-HT transport is highly defective (**Figs. 5D** and **5E**). Thus, both the HA and the HaloTag chimeric proteins report on GCase transport efficiency from the ER to the lysosomes.

Notably, co-precipitation assays reveal that, unlike wild-type GCase (**Fig. 5F**, lane 2, and quantification in **Fig. 5G**) and the GCase_Asn370Ser_ mutant (**Fig. 5F**, lane 3, and quantification in **Fig. 5G**), the GCase_Leu444Pro_ variant associates much stronger with the ER chaperone calnexin (CNX, **Fig. 5F**, upper panel, lane 4, and quantification in **Fig. 5G**) and binding immunoglobulin protein (BiP; **Fig. 5F**, middle panel, lane 4, and quantification in **Fig. 5H**). Thus, the defective delivery to the endolysosomal compartment derives from GCase_Leu444Pro_ variant persistent association with ER-resident chaperones (**Figs. 5F-5H**), indicating the mutant GCase’s inability to attain its native conformation efficiently (Molinari, 2007).

### Halotagged chimeras to monitor defective lysosomal transport of disease-causing GCase mutations by gel electrophoresis

Upon delivery of the chimeric proteins to lysosomal compartments, the HaloTag is cleaved off, generating acid- and protease-resistant fluorescent Halo fragments of about 33 kDa. This allows for direct visualization and quantification of lysosomal delivery of the protein under investigation by monitoring the formation of the 33 kDa fluorescent Halo fragment in gel (Fregno, Fasana *et al*., 2018, Fregno, Fasana *et al*., 2021, Fumagalli, Noack *et al*., 2016, Loi, Raimondi *et al*., 2019, Rudinskiy, Bergmann *et al*., 2022, Rudinskiy and Molinari, 2022, Rudinskiy, Pons-Vizcarra *et al*., 2023).

Cells expressing GCase-HT, GCase_Asn370Ser_-HT or GCase_Leu444Pro_-HT were grown in the presence of 100 nM TMR. Cells were detergent-solubilized, and PNS-separated by SDS-PAGE. TMR-labeled polypeptides were directly visualized in gel with a 532 nm wavelength laser. The PNS of cells expressing GCase-HT contains two fluorescent polypeptides. The upper band corresponds to the 97 kDa full-length GCase-HT (**Fig. 5I**, upper panel, lane 2, FL). The lower band is the 33 kDa Halo fragment, generated upon arrival of the GCase-HT chimera within the lysosomes, where lysosomal enzymes cleave the linker between the GCase protein and the HaloTag (**Fig. 5J**, upper panel, lane 2, HaloTag fragment, quantification in **Fig. 5J**) (Rudinskiy, Bergmann *et al*., 2022, Rudinskiy and Molinari, 2022). The generation of this polypeptide is blocked when cells are exposed to 50 nM BafA1, which inactivates lysosomal enzymes (**Fig. 5I**, upper panel, lane 3). The fluorescent 33 kDa Halo fragment, indicating transport of the chimeric polypeptide to lysosomal compartments, is also generated in cells expressing the GCase_Asn370Ser_ mutant (**Fig. 5I**, upper panel, lane 4, Halo fragment, **Fig. 5J**) but not, or only negligibly, in cells expressing the GCase_Leu444Pro_ mutant (**Fig. 5I**, upper panel, lane 6, Halo fragment, **Fig. 5J**).

Overall, both the imaging (**Figs. 5A‒5E**) and the biochemical assays (**Figs. 5I‒ 5J**) accurately reflect the defective delivery of the GCase_Leu444Pro_ protein, which is associated with severe disease phenotypes linked to *GBA1* mutations.

### GT-02287 and GT-02329 (STARS) restore lysosomal transport of GCase_Leu444Pro_ ***in cellulo***

Next, we quantitatively assessed the performance of GT-02287 and GT-02329 by monitoring variations in the lysosomal generation of the Halo fragment in cells expressing the transport-deficient GCase_Leu444Pro_-HT chimera, using the previously validated quantitative biochemical assay.

Cells expressing GCase_Leu444Pro_-HT were grown in the presence of increasing concentrations of GT-02287 (**Fig. 6A**, lanes 4-7) or GT-02329 (lanes 9-12) for 4 days and exposed to 100 nM TMR for 17h. The cells were detergent-solubilized, and TMR-labeled polypeptides present in the PNS were directly visualized in gel with a 532 nm wavelength laser. Inhibition of lysosomal hydrolases with BafA1 prevents the generation of the Halo fragment (**Fig. 6A**, lane 2 *vs.* lane 3 and **Fig. 6B**, BafA1 *vs.* DMSO). The intensity of the fluorescence of the polypeptide corresponding to the Halo fragment generated in mock-treated cells (**Fig. 6A**, lanes 3, 8 and **Fig. 6B**, DMSO) indicates the basal level of GCase_Leu444Pro_-HT delivered to the lysosomal compartments in these cells, which is much lower than the amount of GCase and GCase_Asn370Ser_ (**Figs. 5I and 5J**). Notably, cell exposure to GT-02287 (**Fig. 6A**, lanes 4-7) or GT-02329 (lanes 9-12) increases the generation of the Halo fragment in a dose-dependent manner (**Fig. 6B**). Thus, both GT-02287 and GT-02329 substantially increase the transport of the GCase_Leu444Pro_ protein from the ER to the lysosome, its site of activity.

**Figure 6.**
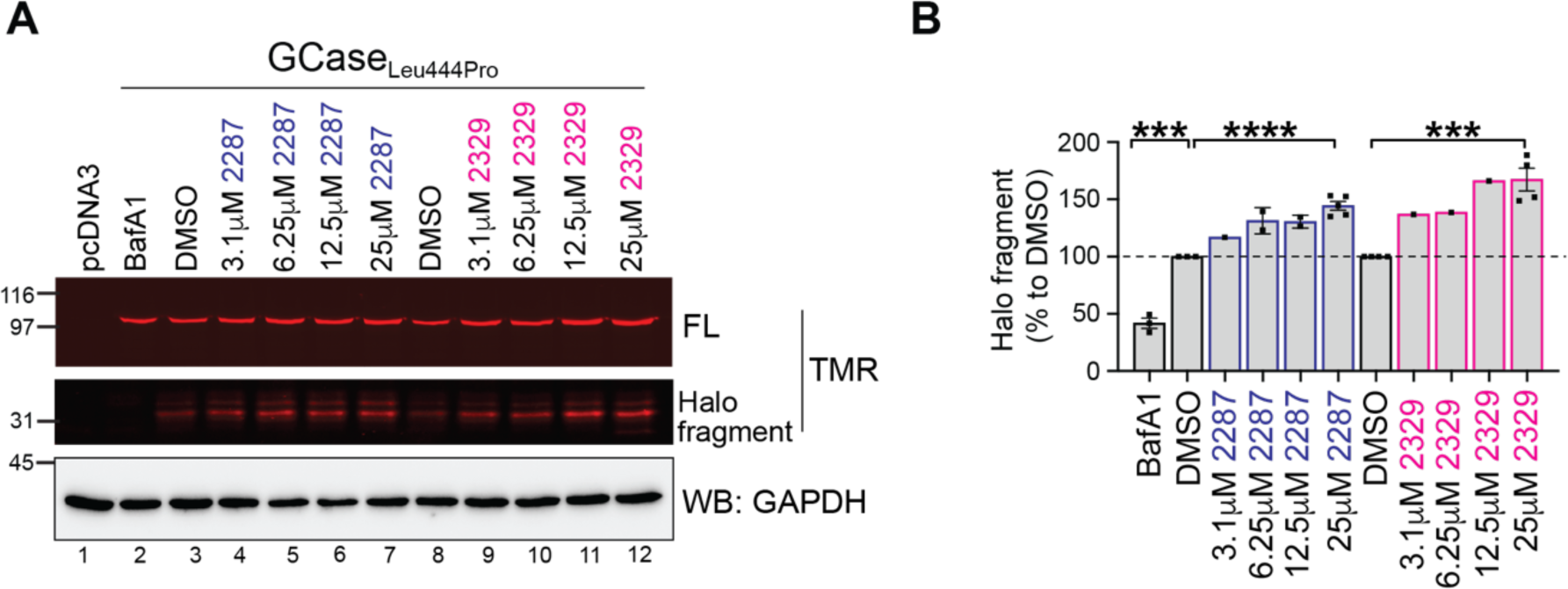
STARS promote lysosomal delivery of the ER-retained variant GCase_Leu444Pro_ and reduce toxic accumulation of GCase endogenous substrate in patient fibroblasts. **(A)** TMR fluorescence of GCase_Leu444Pro_-HT expressed in HEK293 cells treated for 4 days with DMSO or GT compounds (GT-02287 or GT-02329) at concentrations ranging from 3.1 µM to 25 µM. The cells were incubated with 100 nM TMR for the last 17 h. Control cells were treated for 17 h with 50 nM bafilomycin A1 (BafA1; lane 2). After exposure, the proteins were transferred onto a PVDF membrane and immunoblotted against GAPDH. (**B)** Quantification of the Halo fragment, corrected for GAPDH signal and normalized to DMSO. The mean -/+ SEM is shown, with n=4 independent experiments for 25 µM concentrations, n=2 for 6.25 µM and 12.5 µM, and n=3 for BafA1. Statistical analysis was conducted using an unpaired t-test, with ***p<0.001 and ****p<0.0001. TMR, tetramethylrhodamine; HEK293, human embryonic kidney 293 cells; DMSO, dimethyl sulfoxide; BafA1, bafilomycin A1; PVDF, polyvinylidene fluoride; GAPDH, glyceraldehyde 3-phosphate dehydrogenase; SEM, standard error of the mean.

### GT compounds increase intracellular levels of GCase_Leu444Pro_, by promoting its release from ER resident chaperones and by inhibiting its proteasomal clearance from patient-derived primary human fibroblasts

The performance of STARs was assessed in patient fibroblasts containing the homozygous 1448T>C mutation in the *GBA1* gene corresponding to the Leu444Pro substitution. Exposure of the cells to increasing concentrations of GT-02329 for 4 days progressively reduced the co-precipitation of CNX (**Figs. 7A and 7B**) and substantially raised the level of intracellular, endogenous GCase_Leu444Pro_ (**Fig. 7A**, lanes 2-5). An increase in the intracellular GCase_Leu444Pro_ protein in patient cells was also observed upon exposure to a second STAR, GT-02287, as shown by comparing the GCase_Leu444Pro_ level in mock-treated cells (**Fig. 7C**, lane 1) with those exposed to GT-02287 and GT-02329 (lanes 2 and 3, respectively, **Fig. 7D**).

**Figure 7.**
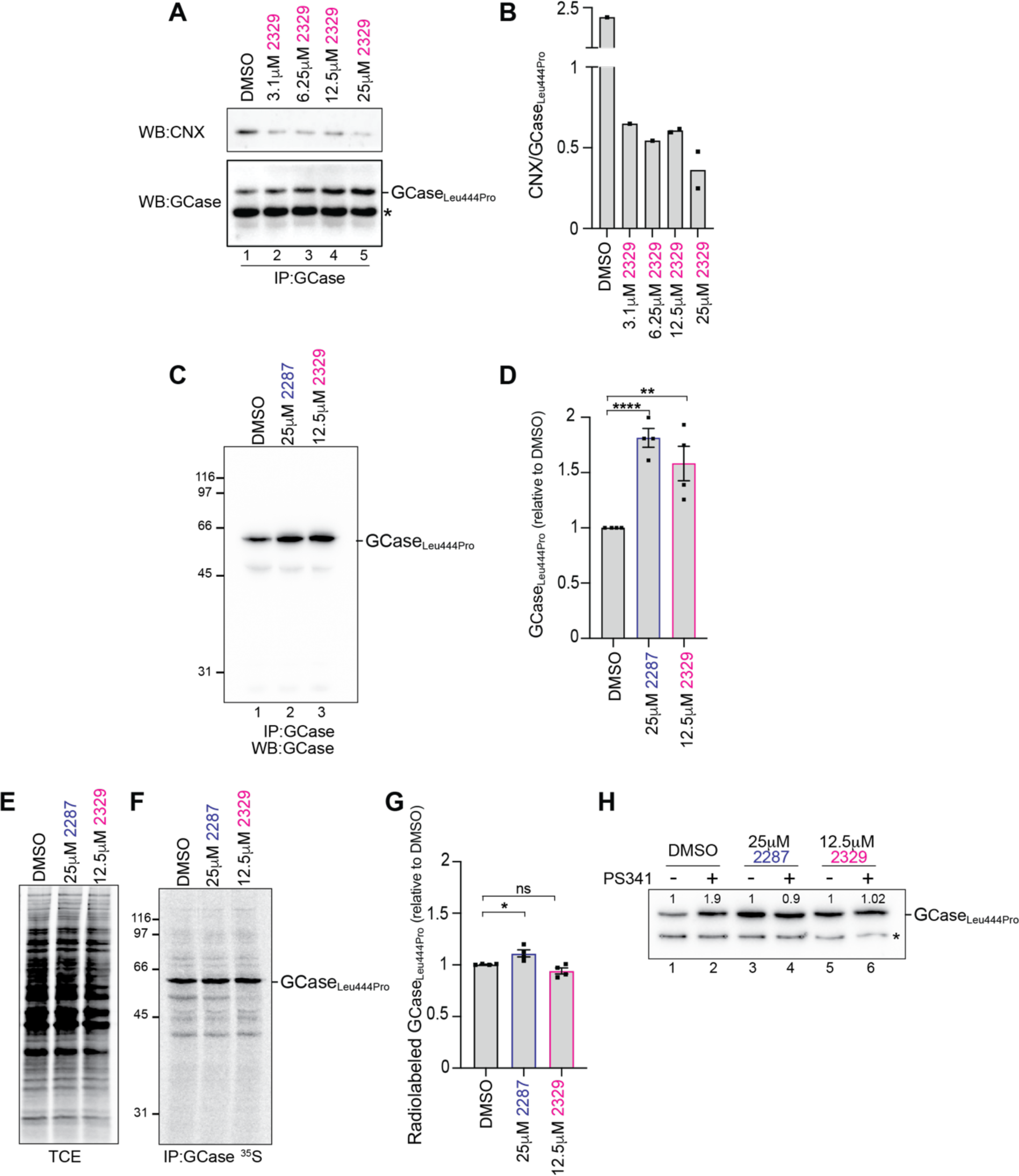
GT compounds promote CNX release and prevent proteasomal degradation of GCase_Leu444Pro_. **(A)** Co-immunoprecipitation with CNX (upper panel) of endogenous GCase_Leu444Pro_ (lower panel) in patient fibroblasts treated for 4 days with DMSO or GT compound GT-02329 at concentrations ranging from 3.1 µM to 25 µM. Protein content in the total cell extracts (TCE) is shown. **\*** unspecific band. (**B)** Quantification of (**A)**. The mean is shown, based on n=2 independent experiments. (**C)** Immunoisolated GCase_Leu444Pro_ from patient fibroblasts treated for 4 days with DMSO or GT compounds (GT-02287 or GT-02329). (**D)** Quantification of (**C)**. The mean ± SEM is shown, based on n=4 independent experiments, using an unpaired t-test with **p<0.01 and ****p<0.0001. (**E)** Radiolabeled protein content in the TCE is shown. (**F)** Radiolabeled GCase_Leu444Pro_ immunoisolate from patient fibroblasts. (**G)** Quantification of (**F)**. The mean ± SEM is shown, based on n=4 independent experiments and using an unpaired *t* test, with ns p>0.05 and *p<0.05. (**H)** GCase_Leu444Pro_ level in patient fibroblasts treated for 4 days with DMSO or GT compounds (GT-02287 or GT-02329) and, for the last 3 h, with the proteasome inhibitor PS341. **\***: unspecific band. CNX, calnexin; DMSO, dimethyl sulfoxide; TCE, total cell extracts; SEM, standard error of the mean; PS341, proteasome inhibitor, bortezomib; ns, not significant.

To assess whether the elevation of GCase_Leu444Pro_ in the patient fibroblasts treated with STARS relies on enhanced synthesis of the endogenous polypeptide, cells were metabolically labeled for 10 minutes with ^35^S-methionine and -cysteine. Analyses of radiolabeled polypeptides (**Fig. 7E**) and radiolabeled GCase_Leu444Pro_ (**Fig. 7F**) in lysates from mock-treated patient fibroblasts (lanes 1) or patient fibroblasts exposed to GT-02287 or GT-02329 (lanes 2 and 3, respectively) show that the STARS do not enhance total protein synthesis or the synthesis of endogenous GCase_Leu444Pro_. Rather, STARs substantially stabilize endogenous GCase_Leu444Pro_ by preventing its proteasomal clearance.

Consistently, in mock-treated cells, the GCase_Leu444Pro_ level is substantially increased when patient cells are exposed to PS341, a selective inhibitor of 26S proteasomes (**Fig. 7H**, lanes 1 vs. 2) (Adams, Palombella *et al*., 1999). Exposure to GT-02287 (lane 3) or GT-02329 (lane 5) raises the level of endogenous GCase_Leu444Pro_ (compare **Fig. 7H**, lane 1 with lanes 3 and 5, **Figs. 7A‒7D**). Incubation with PS341 does not further increase the level, indicating that the turnover of GCase_Leu444Pro_ has already been substantially delayed by the STARS (**Fig. 7H**, lanes 3 vs. 4 and 5 vs. 6).

### GT compounds mitigate UPR in p.L444P/p.L444P patient-derived fibroblasts

The engagement of ER-resident chaperones by folding-defective polypeptides induces low levels of unfolded protein responses, characterized by the transcriptional and translational induction of a very restricted number of ER stress markers (Bergmann, Fregno *et al*., 2018). The persistent engagement of the ER chaperone BiP and CNX by the GCase_Leu444Pro_ mutant (**Fig. 5F**) is mirrored by the constitutively elevated levels of BiP transcripts in p.L444P/p.L444P mutant GD patient fibroblasts as established by real-time qPCR (**Fig. 8A**). This indicates that p.L444P/p.L444P mutant GD patient fibroblasts have higher basal levels of ER stress compared to fibroblasts from healthy patients and from patients expressing the GCase_Asn370Ser_ mutant protein (**Fig. 8A**). Notably, existing literature reports elevated levels of ER stress in mammalian and insect cells expressing mutant forms of GCase (Do, McKinney *et al*., 2019). In sharp contrast, our analyses show that p.N370S/ins patient-derived fibroblasts have normal levels of BiP, consistent with normal ER homeostasis (**Fig. 8A**).

**Figure 8.**
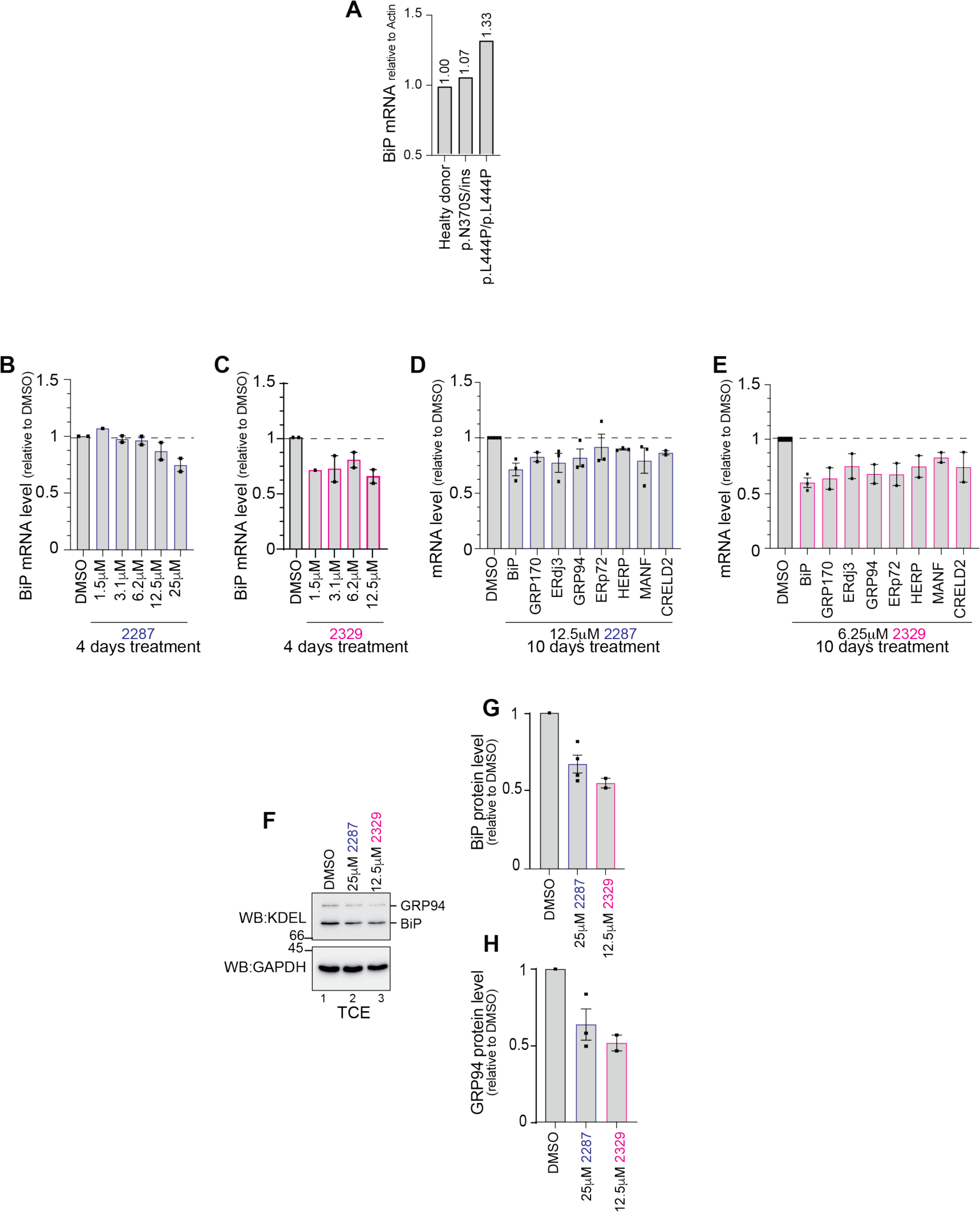
GT compounds alleviate UPR in p.N370S/ins and in p.L444P/p.L444P patient fibroblasts. **(A)** BiP mRNA levels as determined by RT-PCR in healthy donors (lane 1) and patient-derived fibroblasts (lanes 2 and 3). **(B)** BiP mRNA levels monitored by qPCR in GCase_Leu444Pro_ patient fibroblasts (Coriell GM08760) treated for 4 days with DMSO or the STAR GT-02287 at concentrations ranging from 1.5 µM to 25 µM. **(C)** Same as **(B)** for patient fibroblasts treated with GT-02329. **(D)** mRNA levels of selected UPR markers (Bergmann, Fregno *et al*., 2018) monitored by qPCR in GCase_Leu444Pro_ patient fibroblasts (Coriell GM10915) treated for 10 days with DMSO or 12.5 µM GT-02287. **(E)** Same as (D) for cells treated with 6.25 µM GT-02329. **(F)** BiP and GRP94 protein levels monitored by WB analysis in GCaseLeu444Pro patient fibroblasts (Coriell GM08760) treated for 4 days with 25 µM GT-02287 and 12.5 µM GT-02329. (**G** and **H)** Quantification of **(F)**. The mean -/+ SEM is shown. UPR, unfolded protein response; BiP, binding immunoglobulin protein; mRNA, messenger ribonucleic acid; RT-PCR, reverse transcription polymerase chain reaction; qPCR, quantitative polymerase chain reaction; DMSO: dimethyl sulfoxide; STAR, structurally targeted allosteric regulators; GRP94: glucose-regulated protein 94; WB, western blot; SEM, standard error of the mean.

The STARS of GCase substantially reduce chaperone engagement and ER retention of GCase_Leu444Pro_ (**Figs. 8A‒8B**), allowing the delivery of the mutant polypeptide to the lysosomal compartment (**Fig. 6**). This has positive effects by diminishing the lysosomal accumulation of GCase substrates. To verify if this also alleviates the constitutive ER stress found in patient cells, we monitored the transcript and protein levels of the ER stress markers upregulated in cells expressing misfolded polypeptides (Bergmann, Fregno *et al*., 2018). Our experiments show that ER stress levels in patient fibroblasts exposed for 4 days to various concentrations of GT-02287 (**Fig. 8B**) or GT-02329 (**Fig. 8C**), as monitored by quantification of BiP transcripts via real-time qPCR (**Figs. 8B and 8C**) and by monitoring the levels of transcripts of other ER stress markers (**Figs. 8D‒8E**), are reduced by 20-40%. More importantly, the reduction of ER stress is also demonstrated by lower levels of BiP and GRP94 proteins in cells treated with the two STARS of GCase (**Figs. 8F‒8H**).

## Discussion

In the present study, we used the innovative SEE-Tx® drug discovery platform to identify an allosteric druggable binding site on the lysosomal protein GCase. Based on our findings, we designed two STARs of GCase (GT-02287 and GT-02329), which bind the protein outside the catalytic site. GT-02287 and GT-02329 enhance the enzymatic activity of GCase in cell lysates, as well as in primary human fibroblasts from a healthy donor and from GD type I, II and III patients characterized by the expression of the Asn370Ser or Leu444Pro mutant forms of GCase. By reducing the lysosomal accumulation of the GCase’s endogenous substrates, GT-02287 and GT-02329 enhance lysosomal performance, alleviate lysosomal toxicity and reduce cellular stress.

To understand the impact of the disease-causing Asn370Ser and Leu444Pro amino acid substitutions on the fate of the GCase protein, and to characterize the mode of action of the two STARs, we employed a series of experimental readouts based on monitoring the fate of HaloTag chimeras of the proteins under investigation. This approach offers quantitative information on protein trafficking to endolysosomal compartments using biochemical and imaging techniques (Rudinskiy, Bergmann *et al*., 2022, Rudinskiy and Molinari, 2022, Rudinskiy, Pons-Vizcarra *et al*., 2023) and can be implemented with LysoQuant an automated deep learning approach for the quantifying cargo delivery within endolysosomes (Morone, Marazza *et al*., 2020).

We established that the Leu444Pro mutation causes tight ER retention due to engagement with ER-resident molecular chaperones of the lectin (CNX) and the HSP70 families (BiP). The prolonged engagement of these chaperones indicates a significant delay in the mutant polypeptide’s folding process, preventing its release from the biosynthetic compartment and hampering its transport to the lysosomal compartments. Our experiments reveal that the consequences of the Asn370Ser mutation are much milder than those of the Leu444Pro mutation, consistent with the more severe disease progression of patients carrying the latter mutation.

The prolonged engagement of ER chaperones by the GCase_Leu444Pro_ polypeptide eventually triggers a restricted UPR, characterized by the induction of a class of chaperones including BiP, GRP70, ERdj3, GRP94, ERp72, HERP, MANF and CRELD2, recently associated with a transcriptional/translational response to intraluminal accumulation of folding-defective polypeptides (Bergmann, Fregno *et al*., 2018, Bergmann and Molinari, 2018). Notably, GT-02287 and GT-02329 reduce polypeptide retention by the ER chaperone system and protect mutant forms of GCase from unwanted proteasomal degradation, indicating enhanced folding capacity. This has an immediate positive effect, as shown by reduced stress levels in patient cell lines and enhanced delivery of mutant polypeptides to lysosomal compartments.

GT-02287 and GT-02329 have been designed as STARS to bind the target polypeptide outside the active site, as confirmed in our tests by SPR competition experiments with IFG. Consistently, their enhanced delivery to the lysosomal compartments results in the clearance of GCase substrates, whose accumulation interferes with the proper lysosomal functions and is a hallmark of GD. The findings of this study suggest that GT-02287 and GT-02329 have the potential to be developed further as potential therapeutic agents for GCase-related disorders, including GD, PD and Dementia with Lewy Bodies.

In conclusion, our findings indicate that GT-02287 and GT-02329 promote the conformational maturation of disease-causing mutant forms of GCase, facilitate their release from ER-resident molecular chaperones, protect them from unwanted proteasomal clearance, restore lysosomal activity and alleviate misfolded-protein-induced ER stress in cultured cells and in GD patient-derived primary human fibroblasts.

## Supporting information

S1 Table. Antibodies used. (PDF)

S2 Table. Primers for quantitative polymerase chain reactions (qPCR). (PDF)

S1 File. SPR competition with IFG. (PDF)

## Acknowledgments

We thank the members of Molinari’s laboratory for their discussions and critical reading of the manuscript. We also thank Dr Joanne Taylor for her helpful suggestions. Klara J. Belzar, Ph.D., Med-writeplus Ltd., UK, provided medical writing and editorial assistance in preparing the manuscript. Additionally, we appreciate the support and advice on the SPR technique provided by the Technological Centers (CCiTUB) at the University of Barcelona, Barcelona, Spain.

## Author contributions

Conceptualization: NPC, EC, MB, AMGC, MM; Methodology: NPC, AR, AD, EC, AMGC, IF, MR, TS, TJB and MM; Investigation: NPC, AR, AD, EC, IF, MR, TS and TJB; Visualization: NPC, AR, AD, EC, MB, AMGC, IF, MR, TS, TJB and MM; Funding acquisition: MB and MM; Project administration: MB and MM; Supervision: NPC, EC, AMGC and MM; Writing – original draft: NPC, EC, AMGC and MM; Writing – review & editing: NPC, AR, AD, EC, MB, AMGC, IF, MR and MM.

## Data and materials availability

All data and materials used in the analysis are available, without restriction, upon reasonable request to reproduce or extend the analyses.

---

**S1 Table.** Antibodies used in study.

| **Antibody** | **Manufacturer** | **Catalog number** | **Dilution** |
| --- | --- | --- | --- |
| Rat anti-LAMP1 | DSHB | 1D4B | 1:50 (CLSM) |
| Rabbit anti-HA | Sigma | H6908 | 1:100 (CLSM)<br>1:3000 (IB) |
| Mouse anti-KDEL | Stressgen | SPA-827 | 1:1000 (IB) |
| Rabbit anti-CNX | Kind gift from A. Helenius | Not applicable | 1:3000 (IB) |
| Mouse anti-GAPDH | Merck | MAB374 | 1:30000 (IB) |
| Rabbit anti-GCase | Sigma | G4171 | 1:500 (IB) |
| Mouse anti-GCase | Abnova | H00002629-M01 | 1:500 (IB) |
| Protein A HRP-conjugated | Invitrogen | 101023 | 1:20000 (IB) |
| Goat anti-mouse HRP-conjugated | Southern Biotech | 1031-05 | 1:20000 (IB) |
| Goat anti-rabbit AlexaFluor488-conjugated | Thermo Fisher Scientific | A-21206 | 1:300 (CLSM) |
| Goat anti-rat AlexaFluor647-conjugated | Thermo Fisher Scientific | A-21247 | 1:300 (CLSM) |

**S2 Table.**
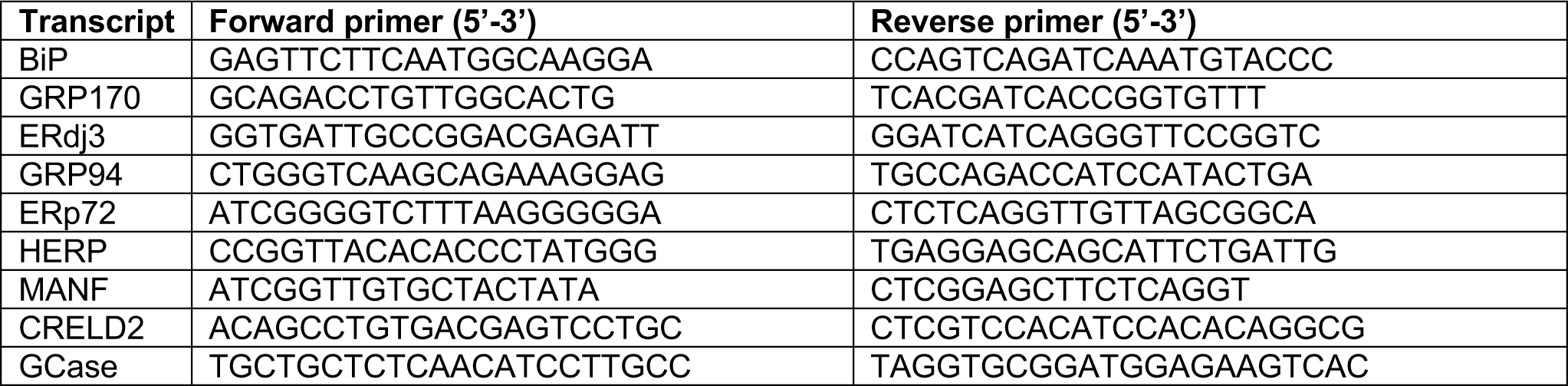
Primers for quantitative polymerase chain reactions (qPCR)

**S1 File. SPR competition with IFG**

Surface plasmon resonance (SPR) competition experiments with isofagomine (IFG) confirmed Gain Therapeutic (GT) compounds bind to Cerezyme at both pHs; however, not at the active site but at a different pocket.

**S1 Fig.**
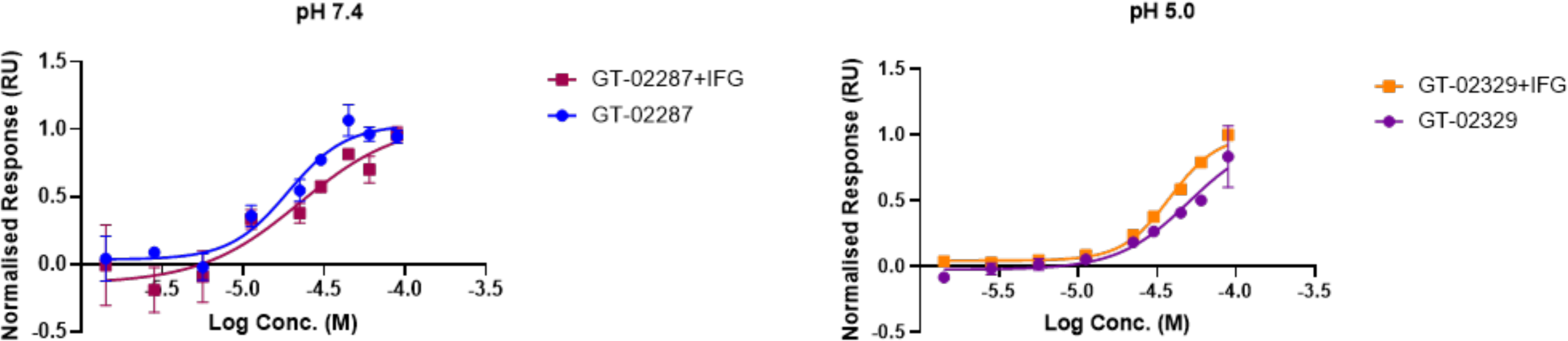
SPR dose-response for GT compounds binding to immobilized GCase monitored at neutral (7.4) and acidic (5.0) pH with and without the presence of the inhibitory chaperone isofagomine (IFG).

## References

Adams, J, Palombella, VJ, Sausville, EA, et al. 1999. Proteasome inhibitors: a novel class of potent and effective antitumor agents. Cancer Res, 59, 2615–2622.

Alonso, XB, Garcia, DA & Schmidtke, P. Patent 2012. Method of binding site and binding energy determination by mixed explicit solvent simulations.

Alvarez-Garcia, D & Barril, X 2014. Molecular simulations with solvent competition quantify water displaceability and provide accurate interaction maps of protein binding sites. J Med Chem, 57, 8530–8539. doi:10.1021/jm5010418.

Alvarez-Garcia, D, Schmidtke, P, Cubero, E, et al. 2022. Extracting Atomic Contributions to Binding Free Energy Using Molecular Dynamics Simulations with Mixed Solvents (MDmix). Curr Drug Discov Technol, 19, 62–68. doi:10.2174/1570163819666211223162829.

Barroso, M, Puchwein-Schwepcke, A, Buettner, L, et al. 2024. Use of the Novel Site-Directed Enzyme Enhancement Therapy (SEE-Tx) Drug Discovery Platform to Identify Pharmacological Chaperones for Glutaric Acidemia Type 1. J Med Chem, 67, 17087–17100. doi:10.1021/acs.jmedchem.4c00292.

Bergmann, TJ, Fregno, I, Fumagalli, F, et al. 2018. Chemical stresses fail to mimic the unfolded protein response resulting from luminal load with unfolded polypeptides. J Biol Chem, 293, 5600–5612. doi:10.1074/jbc.RA117.001484.

Bergmann, TJ & Molinari, M 2018. Three branches to rule them all? UPR signalling in response to chemically versus misfolded proteins-induced ER stress. Biol Cell, 110, 197–204. doi:10.1111/boc.201800029.

Cubero, E, Ruano, A, Delgado, A, et al. 2024. Discovery of allosteric regulators with clinical potential to stabilize alpha-L-iduronidase in mucopolysaccharidosis type I. PLoS One, 19, e0303789. doi:10.1371/journal.pone.0303789.

Do, J, Mckinney, C, Sharma, P, et al. 2019. Glucocerebrosidase and its relevance to Parkinson disease. Mol Neurodegener, 14, 1–16. doi:10.1186/s13024-019-0336-2.

England, CG, Luo, HM & Cai, WB 2015. HaloTag Technology: A Versatile Platform for Biomedical Applications. Bioconjugate Chem, 26, 975–986. doi:10.1021/acs.bioconjchem.5b00191.

Fregno, I, Fasana, E, Bergmann, TJ, et al. 2018. ER-to-lysosome-associated degradation of proteasome-resistant ATZ polymers occurs via receptor-mediated vesicular transport. EMBO J, 37, e99259. doi:10.15252/embj.201899259.

Fregno, I, Fasana, E, Solda, T, et al. 2021. N-glycan processing selects ERAD-resistant misfolded proteins for ER-to-lysosome-associated degradation. EMBO J, 40, e107240. doi:10.15252/embj.2020107240.

Fumagalli, F, Noack, J, Bergmann, TJ, et al. 2016. Translocon component Sec62 acts in endoplasmic reticulum turnover during stress recovery. Nat Cell Biol, 18, 1173–1184. doi:10.1038/ncb3423.

Guzman, B & Taylor, JM 2024. GT-02287, a clinical-stage GCase enhancer, improves activities of daily living and cognitive performance in a preclinical model of GBA1 Parkinson’s disease. Presented at the Federation of European Neuroscience Society Forum. Vienna, Austria.

Kimple, ME, Brill, AL & Pasker, RL 2013. Overview of affinity tags for protein purification. Curr Protoc Protein Sci, 73, 9.9.1-9.9.23. doi:10.1002/0471140864.ps0909s73.

Loi, M, Raimondi, A, Morone, D, et al. 2019. ESCRT-III-driven piecemeal micro-ER-phagy remodels the ER during recovery from ER stress. Nat Commun, 10, 5058. doi:10.1038/s41467-019-12991-z.

Los, GV, Encell, LP, Mcdougall, MG, et al. 2008. HaloTag: a novel protein labeling technology for cell imaging and protein analysis. ACS Chem Biol, 3, 373–382. doi:10.1021/cb800025k.

Maor, G, Rencus-Lazar, S, Filocamo, M, et al. 2013. Unfolded protein response in Gaucher disease: from human to Drosophila. Orphanet J Rare Dis, 8, 1–14. doi:10.1186/1750-1172-8-140.

Molinari, M 2007. N-glycan structure dictates extension of protein folding or onset of disposal. Nat Chem Biol, 3, 313–320.

Morone, D, Marazza, A, Bergmann, TJ, et al. 2020. Deep learning approach for quantification of organelles and misfolded polypeptide delivery within degradative compartments. Mol Biol Cell, 31, 1512–1524. doi:10.1091/mbc.E20-04-0269.

Park, YH, Chen, N, Hu, J, et al. Shire Human Genetic Therapies, Inc. Patent 2019. Formulations comprising glucocerebrosidase and isofagomine.

Patching, SG 2014. Surface plasmon resonance spectroscopy for characterisation of membrane protein-ligand interactions and its potential for drug discovery. Biochim Biophys Acta, 1838, 43–55. doi:10.1016/j.bbamem.2013.04.028.

Premkumar, L, Sawkar, AR, Boldin-Adamsky, S, et al. 2005. X-ray structure of human acid-beta-glucosidase covalently bound to conduritol-B-epoxide. Implications for Gaucher disease. J Biol Chem, 280, 23815–23819. doi:10.1074/jbc.M502799200.

Rudinskiy, M, Bergmann, TJ & Molinari, M 2022. Quantitative and time-resolved monitoring of organelle and protein delivery to the lysosome with a tandem fluorescent Halo-GFP reporter. Mol Biol Cell, 33, ar57. doi:10.1091/mbc.E21-10-0526.

Rudinskiy, M & Molinari, M 2022. Tandem fluorescent Halo-GFP reporter for quantitative and time-resolved monitoring of organelle and protein delivery to lysosomes. Autophagy Rep, 1, 187–191. doi:10.1080/27694127.2022.2061679.

Rudinskiy, M, Pons-Vizcarra, M, Solda, T, et al. 2023. Validation of a highly sensitive HaloTag-based assay to evaluate the potency of a novel class of allosteric beta-Galactosidase correctors. PLoS One, 18, e0294437. doi:10.1371/journal.pone.0294437.

Ruiz-Carmona, S, Alvarez-Garcia, D, Foloppe, N, et al. 2014. rDock: a fast, versatile and open source program for docking ligands to proteins and nucleic acids. PLoS Comput Biol, 10, e1003571. doi:10.1371/journal.pcbi.1003571.

Schindelin, J, Arganda-Carreras, I, Frise, E, et al. 2012. Fiji: an open-source platform for biological-image analysis. Nat Methods, 9, 676–682. doi:10.1038/NMETH.2019.

Schuck, P 1997. Use of surface plasmon resonance to probe the equilibrium and dynamic aspects of interactions between biological macromolecules. Annu Rev Biophys Biomol Struct, 26, 541–566. doi:10.1146/annurev.biophys.26.1.541.

Shaaltiel, Y, Bartfeld, D, Hashmueli, S, et al. 2007. Production of glucocerebrosidase with terminal mannose glycans for enzyme replacement therapy of Gaucher’s disease using a plant cell system. Plant Biotechnol J, 5, 579–590. doi:10.1111/j.1467-7652.2007.00263.x.

